# Human visual cortex and deep convolutional neural network care deeply about object background

**DOI:** 10.1101/2023.04.14.536853

**Authors:** Jessica Loke, Noor Seijdel, Lukas Snoek, Lynn K. A. Sörensen, Ron van de Klundert, Matthew van der Meer, Eva Quispel, Natalie Cappaert, H. Steven Scholte

**Author notes:** Corresponding author (JL). These authors contributed equally / shared first author (Jessica Loke, Noor Seijdel). These authors also contributed equally to this work. The authors declare no competing financial interests. **Conflict of interest:** The authors declare no competing financial interests. **Data and code availability:** Data and code to reproduce the analyses in this article will be made available at https://osf.io/es34u/.

## Abstract

Deep convolutional neural networks (DCNNs) are able to predict brain activity during object categorization tasks, but factors contributing to this predictive power are not fully understood. Our study aimed to investigate the factors contributing to the predictive power of DCNNs in object categorization tasks. We compared the activity of four DCNN architectures with electroencephalography (EEG) recordings obtained from 62 human subjects during an object categorization task. Previous physiological studies on object categorization have highlighted the importance of figure-ground segregation - the ability to distinguish objects from their backgrounds. Therefore, we set out to investigate if figure-ground segregation could explain DCNNs predictive power. Using a stimuli set consisting of identical target objects embedded in different backgrounds, we examined the influence of object background versus object category on both EEG and DCNN activity. Crucially, the recombination of naturalistic objects and experimentally-controlled backgrounds creates a sufficiently challenging and naturalistic task, while allowing us to retain experimental control. Our results showed that early EEG activity (<100ms) and early DCNN layers represent object background rather than object category. We also found that the predictive power of DCNNs on EEG activity is related to processing of object backgrounds, rather than categories. We provided evidence from both trained and untrained (i.e. random weights) DCNNs, showing figure-ground segregation to be a crucial step prior to the learning of object features. These findings suggest that both human visual cortex and DCNNs rely on the segregation of object backgrounds and target objects in order to perform object categorization. Altogether, our study provides new insights into the mechanisms underlying object categorization as we demonstrated that both human visual cortex and DCNNs care deeply about object background.

**Author summary:** Our study aimed to investigate the factors contributing to the predictive power of deep convolutional neural networks (DCNNs) on EEG activity in object recognition tasks. We compared the activity of four DCNN architectures with human neural recordings during an object categorization task. We used a stimuli set consisting of identical target objects embedded in different phase-scrambled backgrounds. The distinction between object backgrounds and object categories allows us to investigate the influence of either factor for human subjects and DCNNs. Surprisingly, we found that both human visual processing and early DCNNs layers dedicate a large proportion of activity to processing object backgrounds instead of object category. Furthermore, this shared ability to make object backgrounds (and not just object category) invariant is largely the reason why DCNNs are predictive of brain dynamics in our experiment. We posit this shared ability to be an important solution for object categorization. Finally, we conclude that DCNNs, like humans, care deeply about object backgrounds.

## Introduction

Deep convolutional neural networks (DCNNs) have entered the computational modeling scene with high predictive performance of both object category and brain dynamics during object categorization tasks (1–4). These predictions on brain dynamics are not limited to low-level image statistics but also include high-level features such as animacy, object category and semantics (5–9). In fact, DCNNs’ predictive performance on visual processes surpassed hand-engineered, biologically-inspired models (e.g. Gabor wavelet filtered, HMAX) because DCNNs are able to achieve high performance on visual tasks (10,11). Traditional mechanistic models generally include few parameters and are tested on simplistic, artificial stimuli such as bar gratings and white noise; in contrast, DCNNs generally include hundreds of thousands to millions of parameters, and are tested on complex and naturalistic stimuli such as photographs of real objects or scenes. But, this claim to fame is not without faults as DCNNs have also been criticized to be black-boxes (12,13) as researchers struggled to understand how millions of parameters work together to perform tasks such as object categorization (14), and also predict brain activity without being trained with brain data (15).

The criticism towards DCNNs become pointed as studies revealed a divergence between humans and DCNNs categorization strategies - humans and DCNNs make mistakes on different images (16–18), DCNNs have an inherent texture bias while humans have an inherent shape bias (19–22), and DCNNs are susceptible to adversarial attacks imperceptible to humans (23,24). While these studies point to differences in categorization strategies, they do not negate the fact that DCNNs can still produce representations which align with human visual processing (25), as reflected in its high predictive performance of brain dynamics. In other words, though certain DCNNs categorization outputs are incorrect, we could probe DCNNs processing stages and find representations which are shared between DCNNs and humans to understand crucial processing steps (7,26). The right question would then be, “which representations are useful and robust for solving the task?”

In this study, we investigated the factors leading to DCNNs’ high predictive power on human visual processing within an object categorization task, focusing on essential representations for solving the task. Prior research has shown the importance of figure-ground segregation (27,28) - the ability to distinguish an image’s foreground and background (i.e. object and background). This ability is especially crucial when the object and its background share similar features such as line orientations, curvatures and colors. Both humans and DCNNs showed enhanced performance when presented with pre-segmented objects compared to objects embedded in backgrounds (29–31). To investigate this further, we used images with identical target objects embedded in varying background complexities, allowing us to isolate human electroencephalography (EEG) recordings and DCNN activity related to target object categorical features versus object background. This approach provides a challenging and naturalistic task while still maintaining experimental control and enables us to identify potentially useful representations in object categorization. Surprisingly, we discovered that large proportions of activity in both human subjects’ EEG recordings and DCNNs’ activity relate to the processing of object backgrounds, rather than object category. Our findings suggest that the ability to distinguish between target object and object background is an essential representation for object categorization.

## Results

In this study, we investigated the factors contributing to the high predictive performance of Deep Convolutional Neural Networks (DCNNs) in human visual processing dynamics. We compared human subjects’ EEG recordings and DCNN activations using Representational Similarity Analysis (RSA; see Materials and methods section). Under the RSA framework, we examined the representations of EEG recordings and DCNN activations using three categorical representational dissimilarity matrices (RDMs; see Materials and methods section) - segmentation, background complexity and object category (see Figure 7). First, we computed partial correlations between the categorical RDMs and EEG RDMs, and between the categorical RDMs and DCNN RDMs. Second, we qualitatively examined the representational structure of DCNNs using t-distributed stochastic neighbor embedding (tSNE; (32). Results from both the partial correlations and tSNE revealed that both EEG recordings and DCNN activations shared a high proportion of activity distinguishing between objects with and objects without backgrounds. Third, to investigate which processing stage (i.e. which layer) was most similar between human subjects and DCNNs, we performed Spearman correlations between EEG RDMs (at every time sample) with DCNN RDMs (per layer). We showed that DCNN layers which correlate highly with EEG recordings are also layers which correlate highly with the categorical RDM of segmentation.

### Object background largely modulates early neural activity in humans

To investigate which of our experimental factors best explained human subjects EEG recordings, we performed partial correlations between the categorical RDMs with EEG RDMs. (See Figure 1) The EEG RDMs correlated highly with segmentation; this correlation had an onset of 86.67ms, W = 79, *p*(Bonferonni corrected) < .01. This was followed by a correlation between the EEG RDMs with background complexity (onset of 90.56ms), W = 197, *p*(Bonferonni corrected) < .01. Finally, there was a much smaller correlation between the EEG RDMs with object category (onset of 110ms), W = 222, *p*(Bonferonni corrected) < .01. The order of onset significance started with segmentation and background complexity, both factors relating to object background, and subsequently arrived at object category. The correlation between the EEG RDMs with segmentation is significantly higher than the correlation between the EEG RDMs with background complexity and object category at ∼87-246ms and ∼343-409ms, *p*(Bonferonni corrected) < .01. The correlation between the EEG RDMs with background complexity is significantly higher than the correlation between the EEG RDMs with object category at ∼87-246ms and ∼343-413ms, *p*(Bonferonni corrected) < .01. Thus, both factors related to object backgrounds have earlier onsets and higher correlations as compared to object category. We can infer three things from these results - 1. object background modulates majority of visual processing signals, not object category, 2. object background modulates visual processing before object category, and 3. the processing of object background begins early (∼87ms) and maintains through ∼409ms.

**Figure 1.**
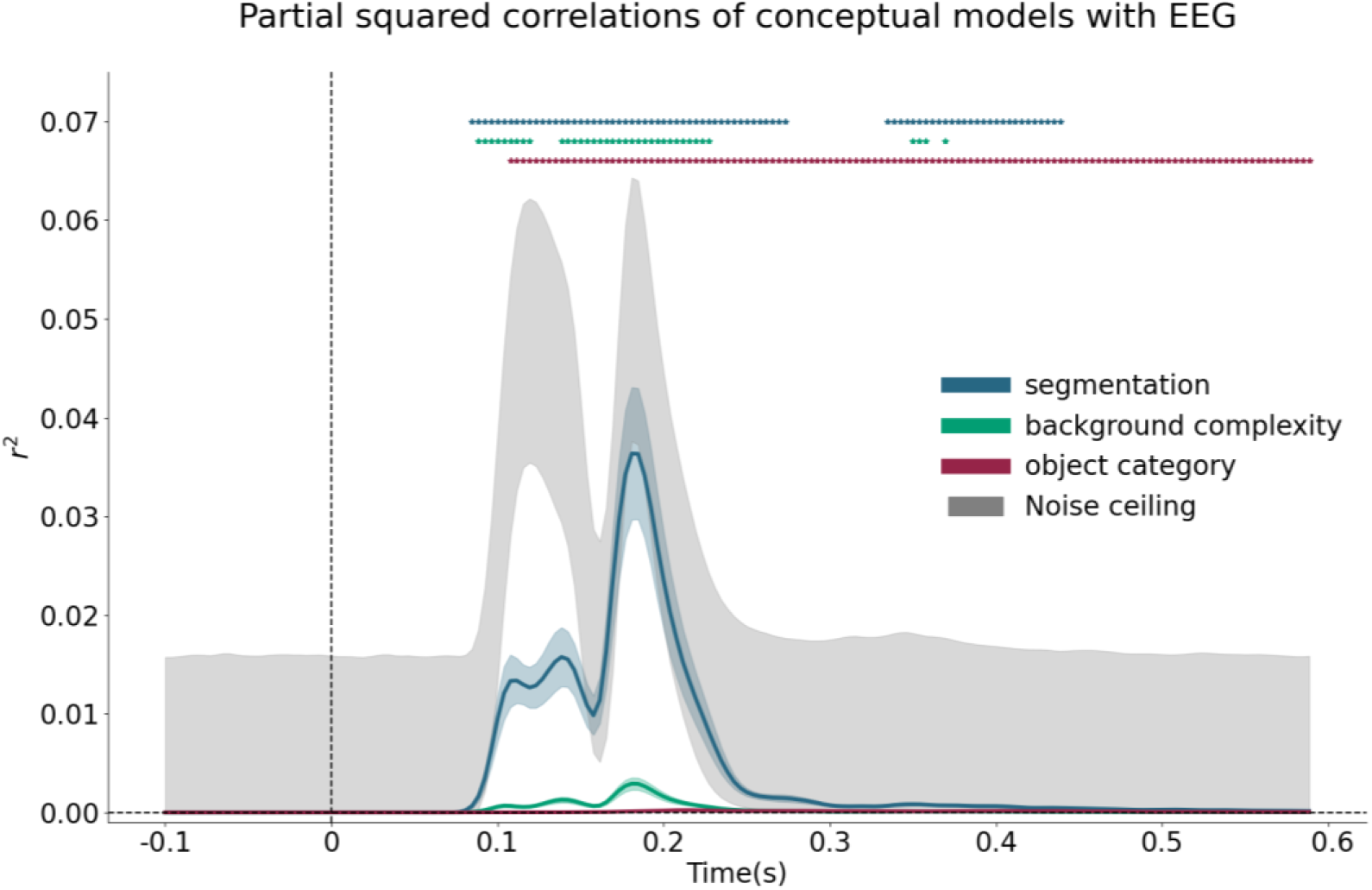
Partial squared correlation of conceptual models with EEG RDMs. By correlating our categorical RDMs with EEG RDMs, we find that the correlation with segmentation was the largest and earliest at 86.67ms. This was followed by the correlation with background complexity with an onset at 90.56ms. Finally, the correlation with object category was much smaller and later at 110ms, compared to both factors related to object backgrounds.

### Object background largely modulates early layers’ activations in DCNNs

Observing that a large proportion of EEG RDMs can be explained by the existence of a background, we similarly performed the partial correlation with DCNNs’ activations, correlating the categorical RDMs with DCNN RDMs (per layer). We have chosen four commonly used DCNNs (AlexNet, VGG-16, ResNet-18, ResNet-50) for predicting brain activity. (See Figure 2) Firstly, we observed that early layers of the DCNNs have high correlation values with segmentation and background complexity - indicating that a large proportion of DCNNs’ early activity was related to object background, not object category, similar to human brains as shown in the previous section. Secondly, we observed that correlations with object category arose in later layers. In deeper networks (with more layers), the correlations with object category became much higher towards the penultimate layer as compared to shallower networks. As a control, we performed the partial correlations between categorical RDMs and untrained DCNN RDMs. We observed that the correlation for segmentation (and not background complexity nor object category) similarly captured a large proportion of untrained DCNNs’ activations. However, unlike their trained counterparts, untrained DCNNs’ correlations arose more gradually and remained until the penultimate (fully-connected) layer. The correlation for background complexity and object category remained close to null throughout the untrained DCNN layers. This indicates that the background differences in untrained DCNNs were not resolved or made invariant, unlike their trained counterparts. Presumably, this transformation of making backgrounds invariant allowed the networks to learn object categorically relevant features.

**Figure 2.**
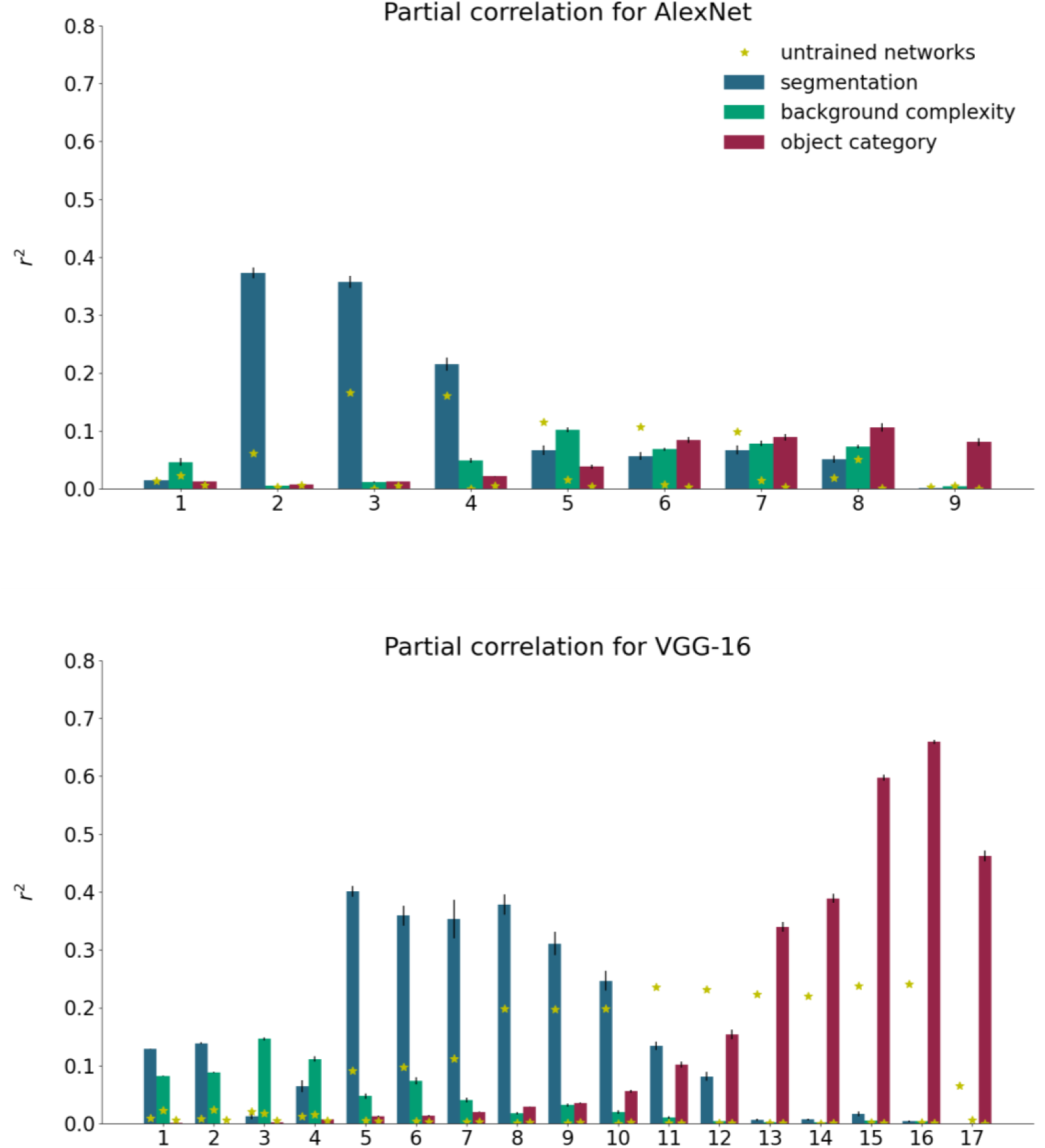
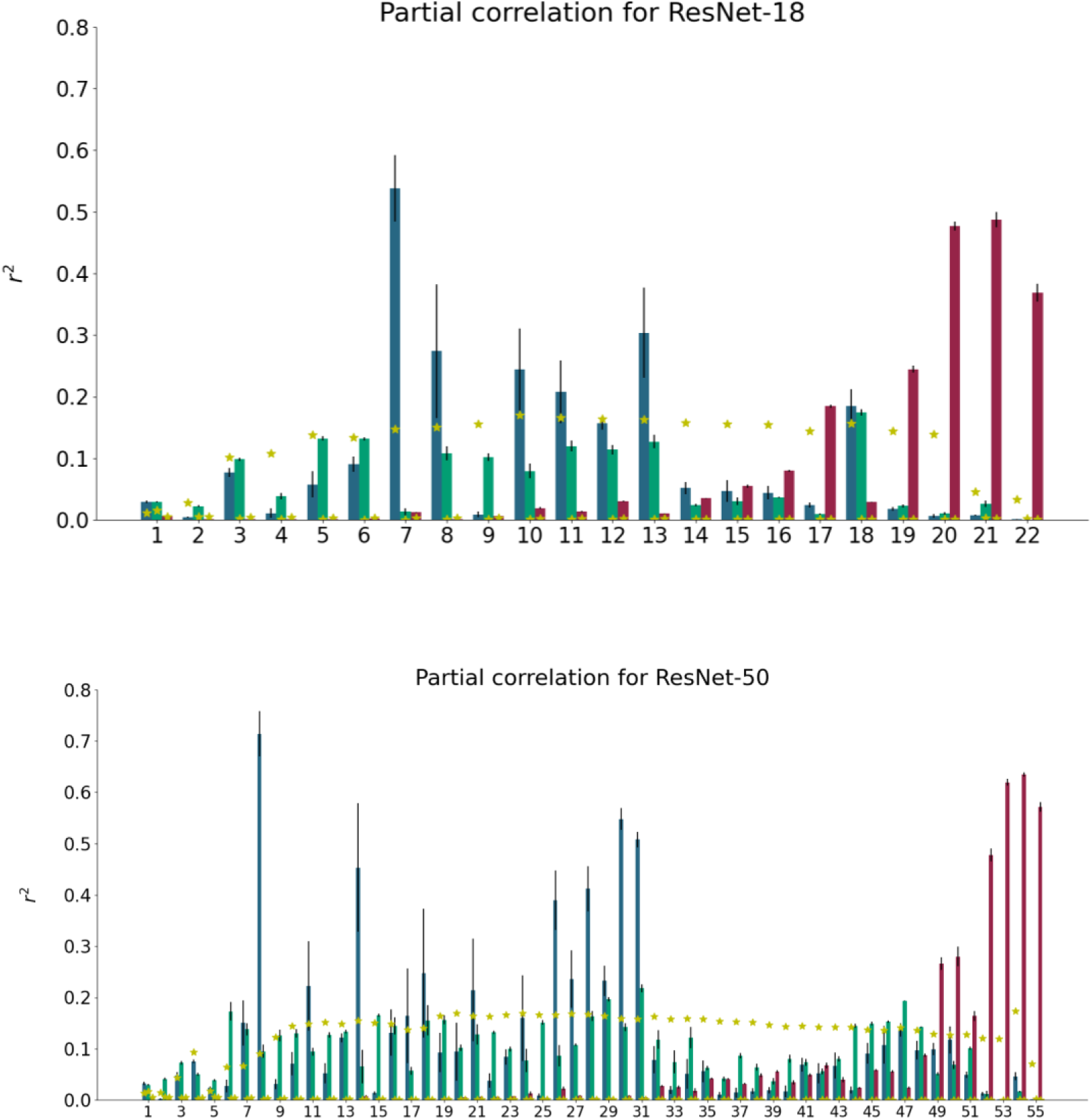
Partial correlation of categorical RDMs with DCNNs. The partial correlations between categorical RDMs (segmentation, background complexity and object category) and DCNN RDMs are shown for each layer of the network. Partial correlations for untrained DCNN RDMs are marked by the yellow stars. Values on the x-axis indicate layer number; values on the y-axis indicate the layer’s partial correlation (in *r²*) with the categorical RDMs. We observed that the early layers of DCNNs correlate largely with both segmentation and background complexity but not with object category. The correlation with object category gradually increases in the later layers, with deeper networks showing a larger increase compared to shallower networks. This pattern of correlation is robust across all networks.

To further understand the network activations, we visualized its activity with t-distributed stochastic neighbor embedding (tSNE; (32)). tSNE maps high-dimensional data points to 2D or 3D spaces. We selected to visualize the activations of DCNNs’ first and final layers, and also the layer with the highest correlation with human subjects EEG recordings. The tSNE visualization showed that with DCNNs layers which correlate most with EEG RDMs, its activation is differentiated along object background - not object category (see Figure 3). In the first layer of all networks, we see a random initialization with no clear clustering of stimuli. In the layer which correlates most with brain activity, we see a clustering of activity according to object backgrounds. And in the final layer, we see a clustering of activity according to object category. With the tSNE visualization, we showed that DCNNs activity differentiates first according to object background and then according to object category. One notable exception of this pattern of results is AlexNet; in its output layer (layer 9), its activity is still clustered along object background. An explanation could be that AlexNet is a much shallower network compared to the other three networks, the lack of depth and additional processing prevents the network from differentiating the stimuli according to their categories.

**Figure 3.**
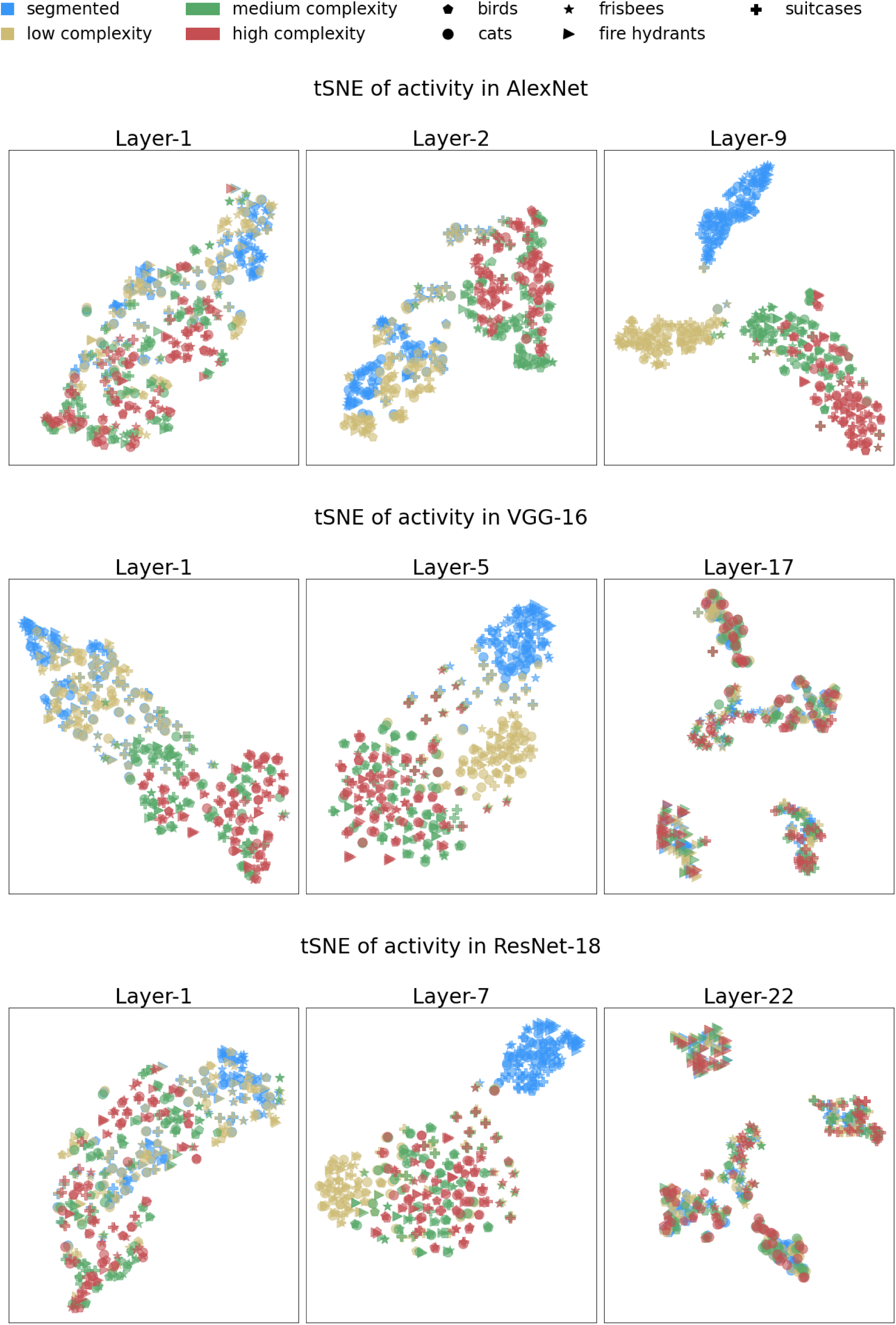
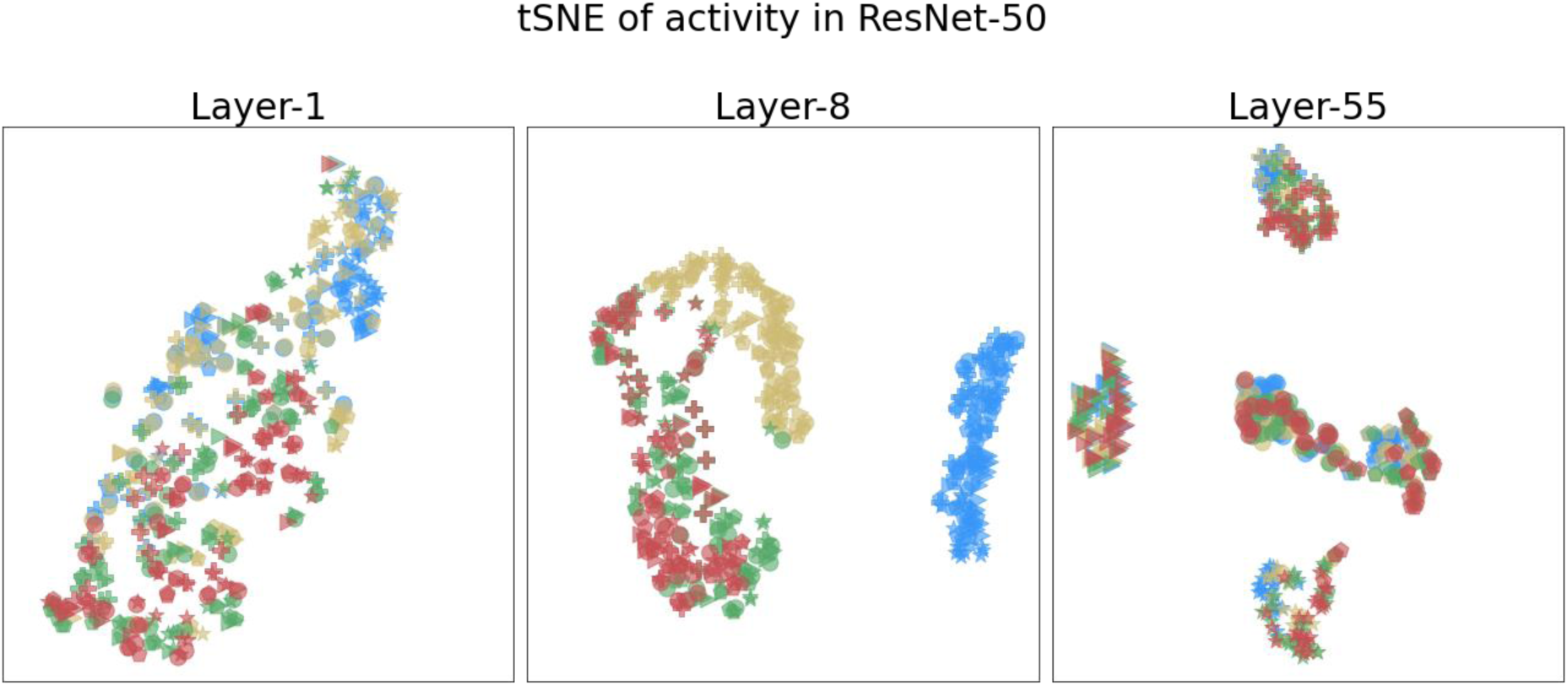
tSNE of DCNNs activations. We applied tSNE to DCNNs’ activations in the first and last layers, and also the layer which correlated most with brain activity. Colors indicate object background conditions - segmented (blue), low complexity (yellow), medium complexity (green), high complexity (red). Markers indicate object category - bird (crosses), cat (circles), frisbee (stars), fire hydrant (triangles), suitcase (plusses). We observed that DCNNs’ activity were differentiated along object background – not object category. In the first layer of all networks, we see a random initialization with no clear clustering of stimuli. In the layer which correlates most with brain activity, we see a clustering of activity according to object background (in colors). In the final layer, we see a clustering of activity according to object category (in marker shapes). Here, we show that DCNNs activity differentiates first according to object background and then according to object category.

As these layers with activations differentiating object background correlate with brain activity, we can infer that DCNNs activity are related to processing object backgrounds. This finding is different from other similar studies using DCNNs because we show that DCNNs layers which capture differences related to object background are also layers which best explain human subject EEG recordings. Additionally, we show that DCNNs layers which capture differences related to object category are also layers which explained the least amount of variance. Thus, both representations from DCNNs and human subjects capture features from object backgrounds, not object category. As such, we posit that the predictive power of DCNNs on brain activity is largely derived from its ability to differentiate object backgrounds, or more specifically, image textures (19).

### Object background predicts brain activity better than DCNNs

Though DCNNs have been touted as the best available mechanistic models, they fell short in explaining human subject EEG recordings as compared to the categorical RDM of segmentation. We have chosen four commonly used DCNNs (AlexNet, VGG-16, ResNet-18, ResNet-50) for predicting brain activity. For each DCNN, we correlated its activation RDMs (per layer) with EEG RDMs (per time sample). (See Figure 4) We observed that AlexNet’s second convolutional layer correlates best with EEG RDMs, followed by VGG-16’s fifth convolutional layer, then ResNet-50’s eighth convolutional layer, and finally ResNet-18’s seventh convolutional layer. Out of the four DCNNs, only AlexNet reached the noise ceiling of the EEG RDMs; whereas, the other networks fell far from the noise ceiling, especially when compared to the categorical RDM of segmentation. We also performed Welch’s t-test between the correlations of DCNNs and EEG, and the correlations of segmentation and EEG, and found that the correlations of DCNNs and EEG significantly differed from the correlations of segmentation and EEG. With the exception of AlexNet conv2 layer - which had higher explained variance as compared to segmentation within the early time window (< ∼160ms), all networks have lower explained variances as compared to segmentation.

**Figure 4.**
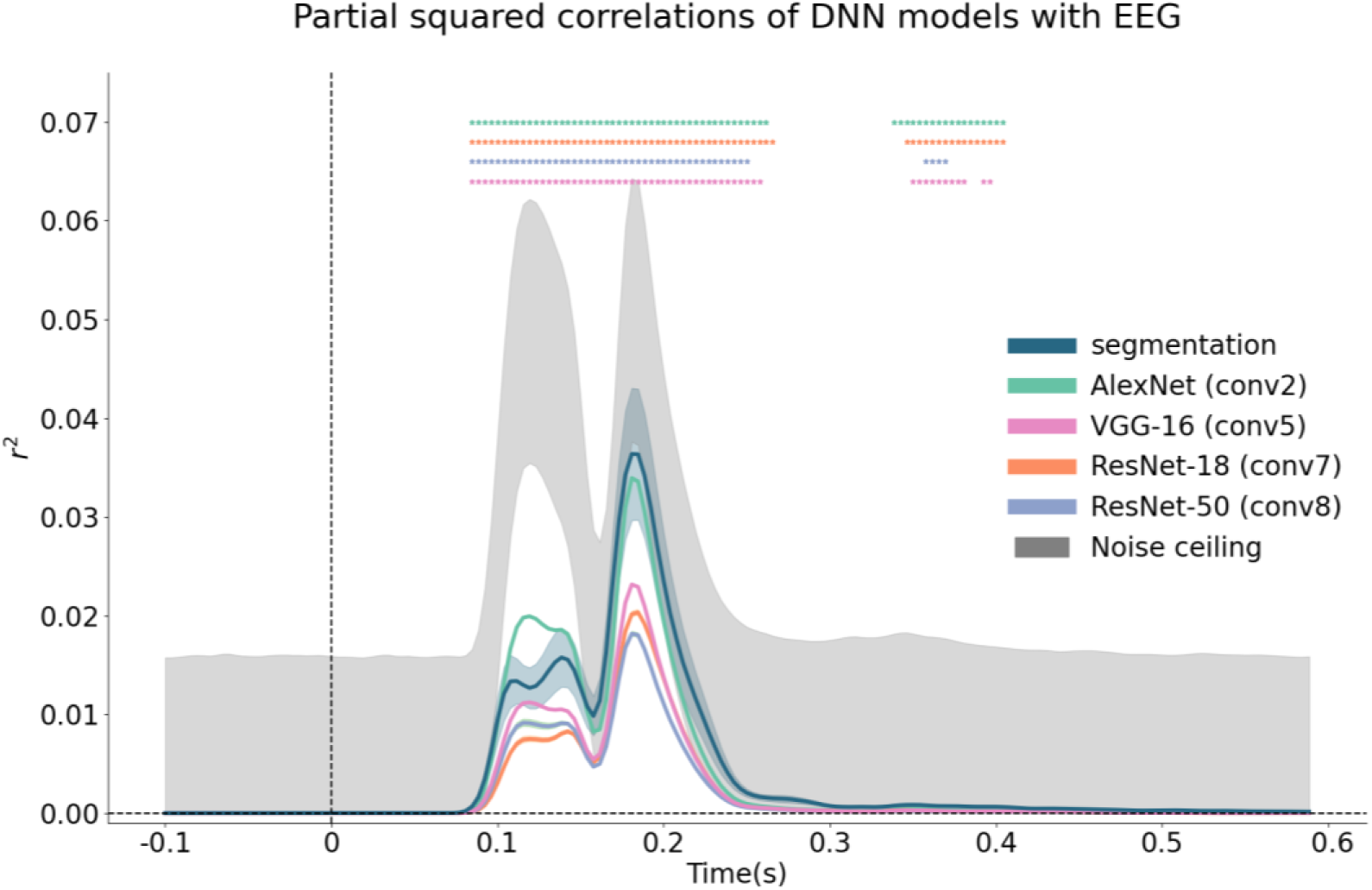
Best correlating DCNNs layers with EEG. We correlated DCNN RDMs (per layer) with EEG RDMs and observed that only AlexNet’s second convolutional layer was close to the noise ceiling of the EEG data. AlexNet was also the only network which surpassed the explained variance of the segmentation model in the earlier time window (< ∼160ms). All other network layers failed to reach the noise ceiling and did not correlate as well with EEG RDMs as compared to the categorical RDM of segmentation.

### DCNNs layers which correlate highly with EEG RDMs also correlate highly with segmentation

After observing that both EEG RDMs and DCNNs RDMs correlate highly with the categorical RDM of segmentation (see Figure 1 and 2), we wanted to investigate the relationship between the RDMs from EEG RDMs, DCNNs RDMs and the categorical RDMs. More specifically, we examined if the correlation values of EEG with a categorical RDM (e.g. segmentation), and the correlation values of DCNNs with the same categorical RDM, correlated with each other. By doing so, we directly investigate if DCNNs’ layers which correlate with a categorical RDM, also correlate well with EEG. This correlation analysis gives us a bridge between EEG and DCNNs to observe if their correlation with a categorical RDM helps explain DCNNs’ predictive power on EEG dynamics. Thus, we took the correlation values of DCNNs with the three categorical RDMs (one datapoint per layer, averaged across five initializations) and plotted its correlation with EEG. We observed that DCNNs RDMs which correlates highly with EEG RDM also correlate highly with the categorical RDM of segmentation (AlexNet, *r=*0.99, *p*<0.01; VGG-16, *r*=0.88, *p*<0.01; ResNet-18, *r*=0.64, *p*<0.01; ResNet-50, *r*=0.62, *p*<0.01). This indicates that DCNNs’ correlation with brain activity is derived from its ability to distinguish between objects’ backgrounds. DCNNs RDMs which correlate highly with background complexity, share a moderate correlation with EEG RDM (AlexNet, *r=-*0.56, *p*=0.11; VGG-16, *r*=0.11, *p*=0.67; ResNet-18, *r*=0.38, *p*=0.08; ResNet-50, *r*=0.27, *p*=0.04). DCNNs RDMs which correlate highly with the categorical RDM of object category actually have a negative correlation with EEG RDMs (AlexNet, *r=-*0.62, *p*=0.08; VGG-16, *r*=-0.72, *p*<0.01; ResNet-18, *r*=-0.35, *p*=0.11; ResNet-50, *r*=-0.12, *p*<0.38).

## Discussion

We set out to investigate the factors leading to DCNNs’ high predictive performance on human visual processing dynamics by studying objects and their backgrounds. Using representational similarity analysis (RSA; (33), we compared the activity of four DCNN architectures with electroencephalography (EEG) recordings of human participants. We focused on three factors: segmentation, background complexity and object category. First, we found that object background largely modulates early EEG signals and early DCNNs layers. Second, we found that both representations from EEG and DCNNs reflected the distinction between objects with and without backgrounds. Third, we showed that the shared distinction of object backgrounds is associated with DCNNs’ high predictive performance on human visual processing dynamics. We posit that DCNNs’ ability to predict EEG signals is derived from its ability to distinguish between target object and object backgrounds.

### Processing of object backgrounds in humans happens earlier and is more substantial than processing of object features

We found high correlations between the categorical RDMs of segmentation and background complexity with EEG - revealing that visual processing (as recorded with EEG) is largely modulated by object backgrounds instead of object category (see Figure 1). Furthermore, the correlations between segmentation and background complexity with EEG have earlier onsets compared to object category - segmentation at 86.67ms, background complexity at 90.56ms, and object category at 110ms. Our result suggests that the processing of object background precedes object features and through this process target objects and their backgrounds becomes distinct. This is evident not only in the latency of significant correlation between the conceptual models and EEG, but also in the correlation between the conceptual models and DCNNs layers - where correlations with segmentation and background complexity precedes object category.

Our finding agrees with previous findings showing that object background complexity influences object categorical perception, with objects embedded in more complex backgrounds to reach categorical perception later (34,35). The longer latency for categorical perception could be explained by time taken to distinguish between the target object and its background. Additionally, our result also extends initial findings that categorical perception is fast (within 150ms) (36,37). Results from earlier studies demonstrating the quickness of categorical perception holds when the presented stimuli was simple (i.e. object with a plain background); however, if the presented stimuli was more complex (i.e. object with a complex background), longer latency incorporating additional processing steps would be required (38). As natural scenes comprises a myriad of complexities in backgrounds, we recommend a careful consideration of not only object category but also backgrounds.

### DCNNs processes on object backgrounds are explaining EEG activity

In our experiment, we show that DCNNs predictive power on EEG data is derived from DCNNs’ inherent ability to distinguish between objects with and without backgrounds. Crucially, the distinction of object backgrounds is orthogonal to the object categorization task. The selected DCNNs for the experimental task have been pre-trained on a naturalistic dataset (ImageNet), and further optimized with a separate dataset (MSCOCO). Nonetheless, DCNNs activations reflect a distinction between objects with and without backgrounds. The distinction is apparent in its partial correlation with the categorical RDMs of segmentation and background complexity (see Figure 2), especially in DCNNs early and mid-layers. Additionally, we also showed that DCNNs layers which correlated with segmentation also correlated with EEG (see Figure 5), suggesting that DCNNs’ predictive power on EEG data is largely derived from the shared ability of both modalities to distinguish between the target object and its background.

**Figure 5.**
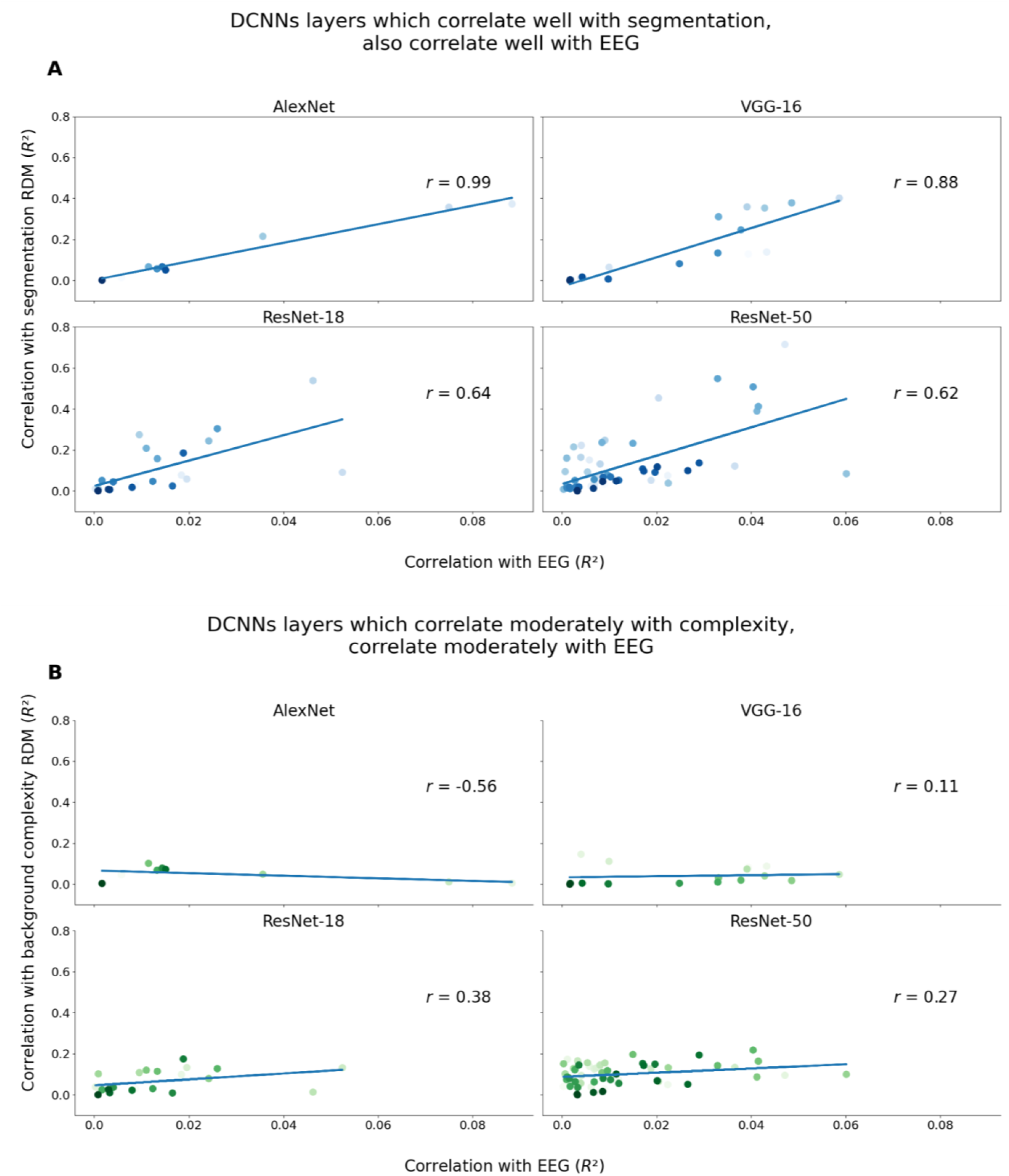
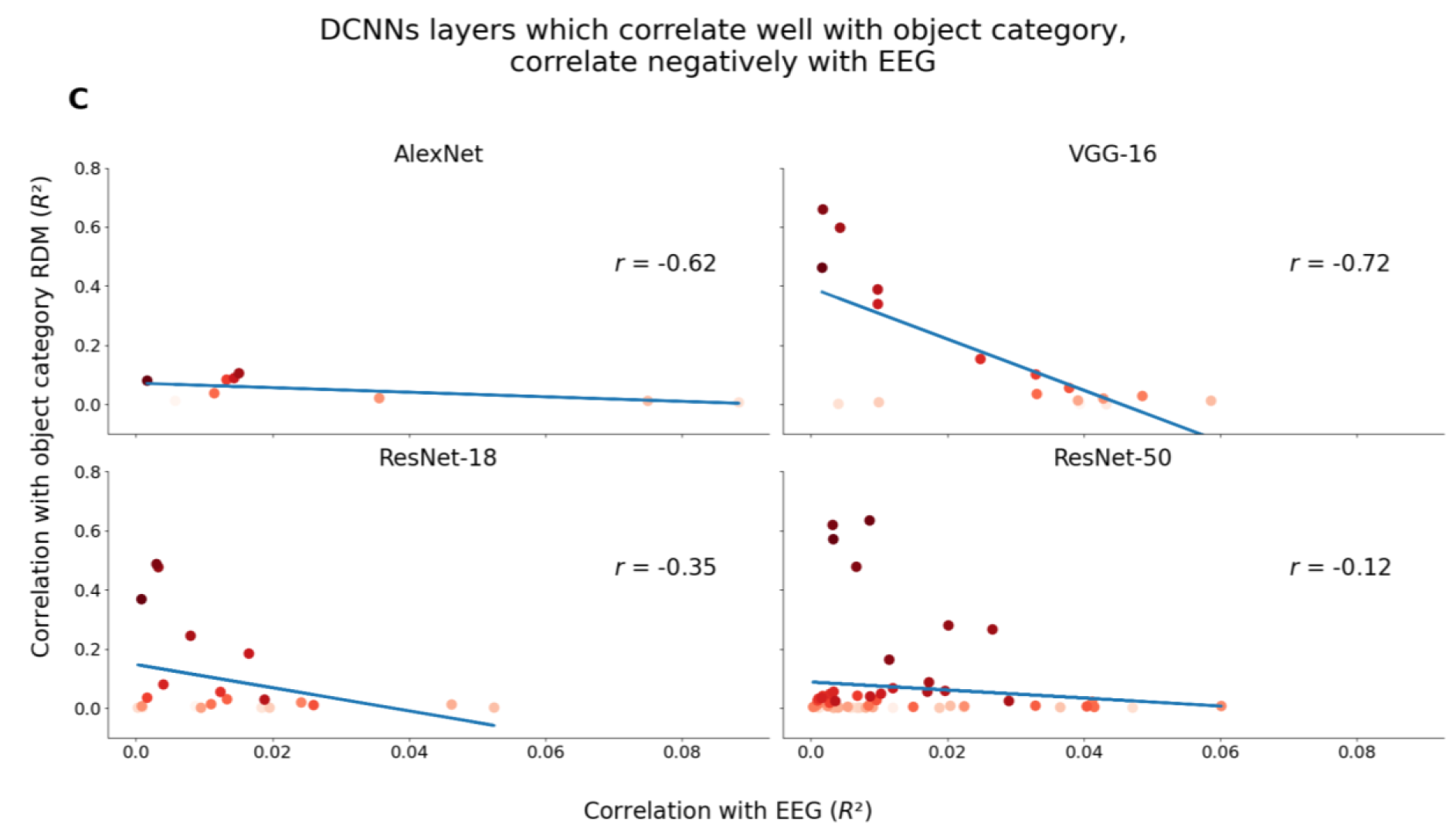
Relationship between DCNNs correlation with EEG and categorical RDMs. Each dot represents a DCNN layer (averaged across five initializations). Darker colors indicate deeper layers within a network and lighter colors indicate shallower layers. A) We observed that layers which correlate highly with EEG are also layers which correlate with the categorical RDM of segmentation. B) There is a moderate relationship between DCNNs’ correlation with EEG and the categorical RDM of background complexity; and C) a negative correlation between DCNNs’ correlation with EEG and the categorical RDM of object category - indicating that DCNN layers which correlate highly with object category actually become dissimilar with EEG RDMs.

Our conclusion that DCNNs’ predictive power on EEG data is derived from the shared ability of both modalities to distinguish between objects’ backgrounds needs to be considered carefully because we have reconstructed an experimental dataset with target objects embedded within artificial backgrounds. There is a high necessity to identify the target object as separate from its background because the artificial backgrounds are uninformative on the object category. In contrast, if the object category correlated with its background (e.g. frisbee with the background of a park), and if the discrimination of object categories could be performed sufficiently well based on the object backgrounds, no distinction needs to be made between target objects and their backgrounds. In reality, most naturalistic scenes will have backgrounds which are informative of its target objects’ categories as these are a matter of statistical correlations. In our study, we constructed an object categorization task which required the distinction of target object and its background with the intention of investigating the mechanism of figure-ground segmentation; surprisingly, we found that both DCNNs and our human subjects shared this ability.

### Emergence of shared solutions for object categorization

The shared ability to distinguish between target objects and their backgrounds within human visual processing and DCNNs affords us to ask a follow up question - “Why does it exist?” This ability was not directly implemented in both systems yet emerged as part of the solution for categorizing objects. Within vision neuroscience, this ability to distinguish between target objects and its backgrounds has long been studied as part of processes known as perceptual grouping or figure-ground segmentation (27,28,39–42). Specifically, these processes refer to the grouping of image elements which belong to different entities. It has been shown that if these processes were interrupted in human subjects, object categorization becomes impaired (43). In our study, the emergence of a shared solution (i.e. perceptual grouping) for object categorization suggests it to be a crucial solution for the task at hand and could elucidate the evolutionary constraints on the problem (44). This helps us arbitrate which biological processes are necessary to incorporate in artificial systems depending on their contexts.

### Figure-ground segregation assists object features learning

Previous research has shown the surprising prediction performance of random weights networks (26,45,46); it is indeed impressive that random weights networks are able to explain any brain activity at all. Our experimental results similarly showed that untrained networks can explain variance in brain activity through its inherent ability to process low-level image statistics. Through correlating untrained networks RDMs with conceptual RDMs, we find that the networks’ activity is modulated only by object background and not object category at all (see Figure 2). We observed a similar predictive performance of an untrained network on V1 in previous studies, where the correlation of the untrained network gradually increased in the early layers and remained until the late layers (46). In our study, we observed that the conceptual RDMs of segmentation correlated highly with the layers of untrained networks, whereas, the conceptual RDMs of background complexity and object category did not correlate with the layers of untrained networks. This indicates that untrained networks are able to distinguish between objects with and without backgrounds, but are unable to distinguish between the background types or categorical features. In contrast, layers of trained networks show a correlation with segmentation up until the middle layers of the network which then gradually decreased, matched by the gradual increase of correlation with object category. This suggests that trained networks “resolved” figure-ground segregation, allowing it to learn object categorical features.

## Conclusion

In summary, we have tested the best mechanistic models of visual processing and showed that both early human visual processing and early DCNN layers are highly modulated by object background, not object category. Moreover, the shared ability to distinguish between object backgrounds explains DCNNs’ predictive power on EEG activity. Neither humans nor DCNNs were explicitly taught to distinguish between object backgrounds but the shared solution emerged to resolve the experimental task of object categorization. Altogether, we have shown that both human visual processing and DCNN care deeply about the object backgrounds.

## Materials and methods

### Data

The electrophysiological data are from (35), it consists of electroencephalography (EEG) recordings from human subjects (*n*=62, 18-35 years old). For a brief description of the experimental paradigm and example of stimuli, please see Figure 6.

**Figure 6.**
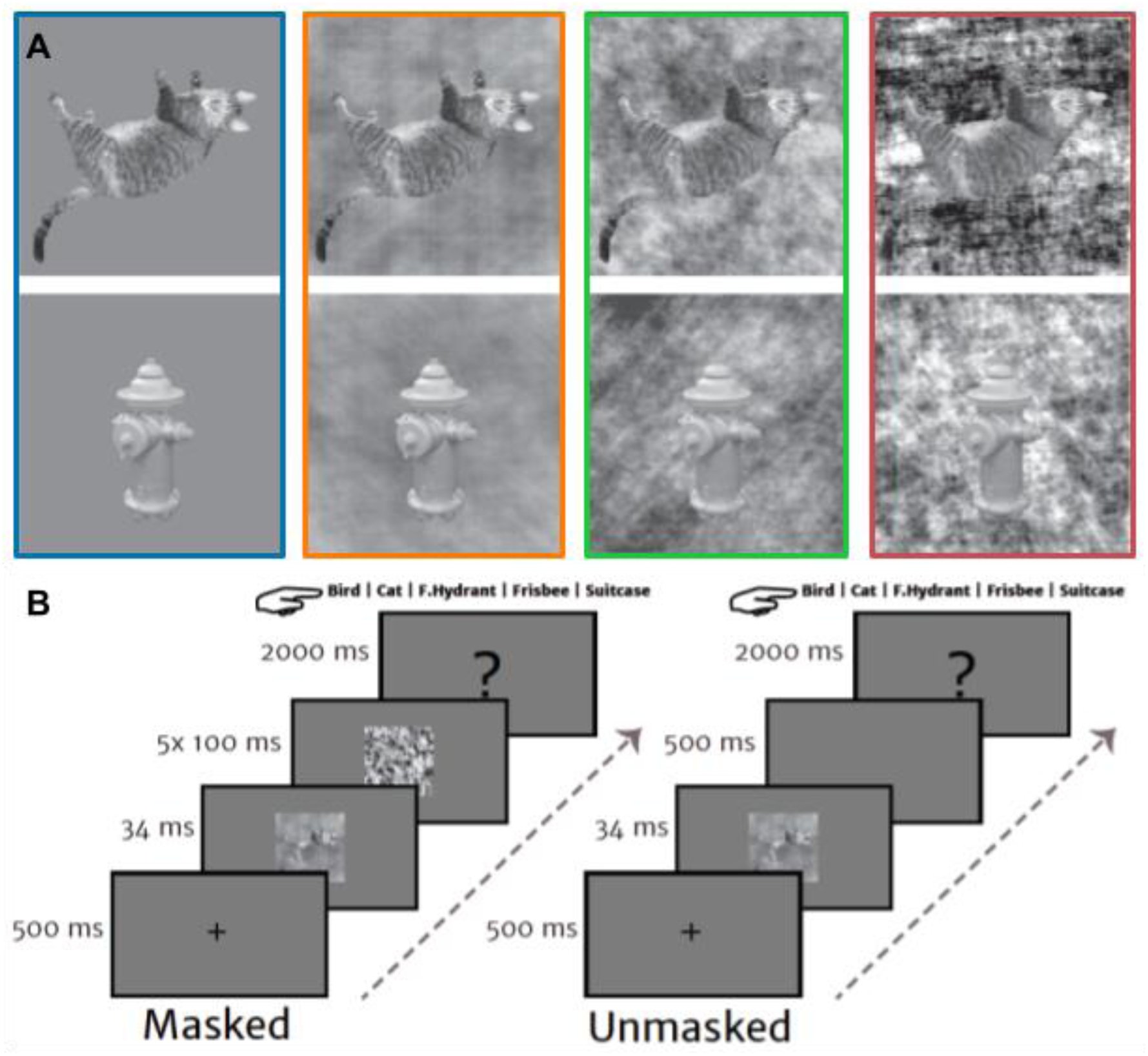
Stimuli sample and experimental paradigm. A) Two object exemplars (cat and fire hydrant) are displayed across four background types. The first (highlighted in blue) is a uniform gray background, referred to as the “segmented” condition. The second (highlighted in orange), third (highlighted in green) and fourth (highlighted in red) are a low, medium and high complexity background respectively.. The increasing levels of background complexity makes it increasingly difficult to differentiate the target object from its background. B) The experimental paradigm had human subjects perform an object categorization task. Each trial starts with a fixation cross of 500ms, followed by a stimulus presentation of 34ms. For masked trials, stimulus presentation is followed by five visual masks, each presented for 100ms. For unmasked trials, the stimulus presentation is followed by a blank screen for 500ms. Finally, there is a response screen displaying the five object category options for 2000ms. Participants completed a total of 960 trials - 120 trials per image condition both masked and unmasked. In this paper, only the unmasked trials were used as our study did not pertain to a comparison of feedforward versus feedback processing. Figure taken from (35).

**Figure 7.**
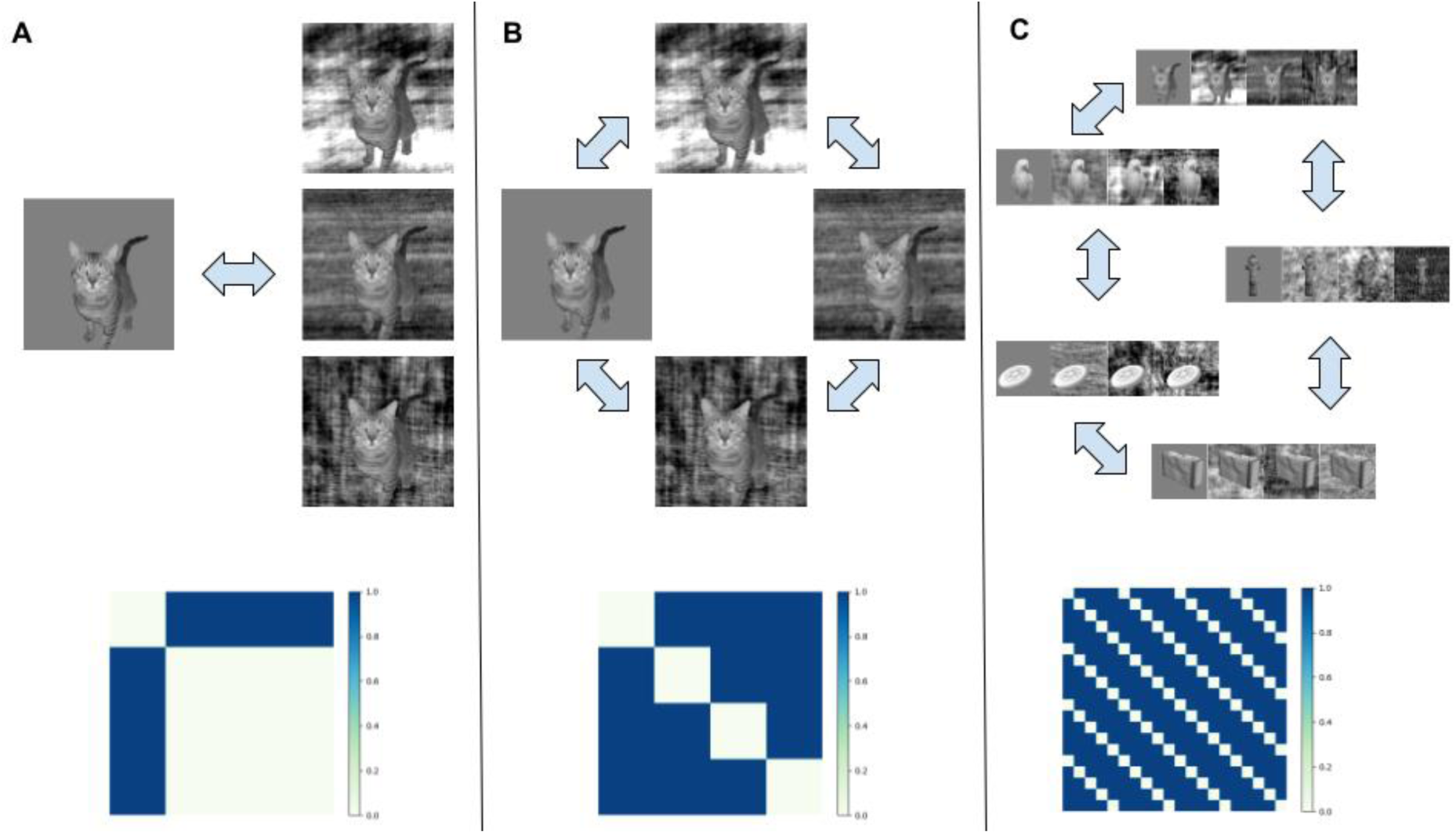
Categorical models of main experimental manipulations. A) The categorical RDM of segmentation distinguishes between trials with and without backgrounds. B) The categorical RDM of background complexity distinguishes between trials with different background complexities. C) The categorical RDM of object category distinguishes between trials based on the target object category.

### Stimuli

The stimuli used consisted of 120 unique target objects (24 per category) from five categories (bird, cat, fire hydrant, frisbee, and suitcase), embedded within four background types (uniform gray background, low complexity, medium complexity and high complexity). This gave us a total of 480 unique stimuli. The backgrounds were created by phase-scrambling the original image backgrounds to remove information aiding recognition of the target object. The complexity of these phase-scrambled backgrounds varied with contrast, with higher contrast indicating higher complexity. The segmented condition does not have phase-scrambled backgrounds but a uniform gray one. The stimuli were presented at a resolution of 512 x 512 pixels.

### Deep convolutional neural networks (DCNNs)

We selected four established DCNN architectures, commonly used in computational modeling - AlexNet (47), VGG-16 (48), ResNet-18 and ResNet-50 (49). Five different seeds of each network were initialized and trained with the ImageNet Large Scale Visual Recognition Challenge 2012 (ILSVRC) dataset, then fine-tuned to the experimental object categories with the Microsoft COCO dataset (50). We used different seeds to capture variance between different initializations and obtain reliable results (51). For the initial training on ILSVRC, we used a learning rate of 0.1 (except for VGG-16 which needed a lower learning rate of 0.05) with a learning rate decay of 0.1 every 30 epochs and a weight decay of 1e-4. We also used a stochastic gradient optimizer with a momentum of 0.9. AlexNet, ResNet-18 and ResNet-50 were trained for 150 epochs while VGG-16 was trained for 74 epochs. All DCNNs reached similar performance accuracies reported in the original papers. For fine-tuning, we replaced the last fully-connected layer and retrained weights from all layers. We fine-tuned the network with a learning rate of 1e-3 with a learning rate decay of 0.1 every 7 epochs. The fine-tuning was performed for 20 epochs. We also used a stochastic gradient descent optimizer with a momentum of 0.9 for fine-tuning. In addition to trained networks, we initialized five different seeds of each architecture with no training as untrained networks. All DCNNs training and fine-tuning was done in PyTorch (52).

### Analysis: Representational Similarity Analysis (RSA)

We used the framework of Representational Similarity Analysis (RSA; (33) to compare EEG activity with DCNNs activations. RSA is a method of analysis allowing for the comparison between different modalities by first generating a representational structure of the stimuli set as reflected in brain activity (as recorded using EEG sensors) and DCNNs (as reflected through its unit activations), and then comparing both those representational structures. This abstraction from EEG sensors and DCNNs unit activations allows us to compare the transformations performed by both modalities on the stimuli. Using RSA, we obtained time-resolved EEG activity and layerwise DCNN activations in the form of representational dissimilarity matrices (RDMs). The RDMs consist of pairwise distances computed from multivariate responses (i.e. pattern of EEG activity or pattern of layerwise DCNNs activations) towards every possible stimuli pair. Pairwise distances were computed as (*1* − *pearson correlation*). An entry in the RDM between stimuli A and B would be - *1* − *pearson correlation of multivariate responses towards stimuli a and B*; whereas, an entry in the RDM between stimuli A and A would be 0. With 480 unique stimuli (120 unique objects x 4 background types), we obtained 480x480 RDMs. In all analyses using RDMs, we used only the upper triangle (excluding the diagonal) since the RDMs are symmetrical.

RDMs of EEG recordings were computed using 22 posterior electrodes (Iz, I1, I2, Oz, O1, O2, POz, PO3, PO4, PO7, PO8, Pz, P1, P2, P3, P4, P5, P6, P7, P8, P9, P10). These electrodes are chosen to focus on activity from visual processing areas and were confirmed in previous studies (34,35). The electrodes placement followed a 10-10 layout, modified with two additional occipital electrodes (I1 and I2) replacing two frontal electrodes (F5 and F6). RDMs were computed from every time sample from -100ms to 600ms relative to stimulus onset. RDMs of DCNNs activations were obtained from activity of all convolutional, pooling and fully-connected layers.

In addition to RDMs from EEG and DCNNs, we also constructed categorical RDMs to evaluate the main effects of our experimental manipulations. We built three categorical RDMs - segmentation, background complexity and object category (see Figure 7). All three RDMs consisted of binary values: “0” representing pairs from the same group, and “1” representing pairs from different groups. Segmentation distinguishes between stimuli with and without backgrounds (see Figure 7A). Background complexity distinguishes between the four background types (see Figure 7B): segmented (no background), low complexity, medium complexity and high complexity. Object category distinguishes between the five object categories (see Figure 7C). Here, it should be noted that the categorical RDMs of segmentation and background complexity correlate substantially (*r* = .45), because the segmented stimuli all have the same complexity (i.e., 0; see Figure 7A & B). As such, to separate the variance associated with segmentation or background complexity, we performed partial correlations between the categorical RDMs and EEG RDMs.

First, we performed partial correlations between the categorical RDMs and EEG RDMs, and between the categorical RDMs and DCNN RDMs to identify the shared representational structure. We chose to use a partial correlation instead of a regression to control for the correlation between the segmentation and background complexity categorical model. Second, we qualitatively inspected the representations from both EEG and DCNNs using t-distributed stochastic neighbor embedding (tSNE) (32). Third, we performed a Spearman correlation (i.e. classical representational similarity analysis) between EEG RDMs (for every time sample) and DCNN RDMs (per layer). Fourth, we normalized each layer’s explained variance from the Spearman correlation against the upper noise ceiling (the upper bound of EEG data) for all time samples and then plotted its median correlation against the layer’s correlation with the categorical RDMs. This allowed us to summarize each layer’s correlation with EEG data across all time samples.

All statistical analysis was performed and visualized in Python using the following packages: NumPy, SciPy, Statsmodels, Pandas, Seaborn, Matplotlib (53–58).

### Analysis: Statistical

We used a Wilcoxon signed rank test to determine the onset of correlation significance between categorical RDMs and EEG RDMs, and to determine statistical significant differences in the correlation values of categorical RDMs. The *p*-values obtained from the Wilcoxon signed rank test are Bonferroni corrected for multiple comparisons (α=0.01).

## Acknowledgements

This work is supported by an Interdisciplinary Doctorate Agreement from the University of Amsterdam to H. Steven Scholte and Natalie Cappaert and an Advanced Investigator Grant from the European Research Council (ERC) to Edward de Haan (#339374).

